# *In vivo* acquired resistance to tamoxifen is associated with irreversible loss of IGF1R, upregulation of insulin receptor and enhanced sensitivity to insulin

**DOI:** 10.64898/2026.06.08.727558

**Authors:** Katelyn J. Hoff, Priya Periakaruppan, Courtney S. Baar, Valeria C. Torres Irizarry, Deepali Sachdev

## Abstract

Selective estrogen receptor modulators (SERMs) like tamoxifen are used to treat hormone receptor positive (HR+) breast cancers. Several models of *in vitro* acquired resistance to tamoxifen using HR+ breast cancer cell lines have been developed. There are few models of resistance to tamoxifen that are generated *in vivo*. Here, we generated a model of *in vivo* resistance to tamoxifen. We show that in vivo tamoxifen resistance is associated with loss of IGF1R. This loss is irreversible as it persists even when tamoxifen is removed. *In vivo* tamoxifen resistance is also associated with upregulation of insulin receptor, specifically the A isoform of IR (IR-A). These resistant cells show enhanced insulin sensitivity compared to the parental endocrine sensitive or *in vitro* acquired tamoxifen resistant cells. Finally, we demonstrate that *in vivo* resistance to tamoxifen results in enhanced proliferation in response to insulin. These studies establish that some mechanisms of *in vivo* resistance to tamoxifen differ from those observed for *in vitro* acquired resistance and increased dependence on insulin signaling drives endocrine resistance in HR+ breast cancer. Taken together, these studies present that targeting IR-A in combination with newer targets of tamoxifen resistance could be tested in endocrine resistant breast cancer.

## Introduction

Breast cancer is the second leading cause of mortality in females (1). 70 to 75% of breast cancers diagnosed express estrogen receptor (ER) and are classified as hormone receptor positive (HR+) breast cancer (2). These patients are treated with endocrine therapy (ET) that target the function or expression of ER (3). Selective estrogen receptor modulators (SERMs) such as tamoxifen, aromatase inhibitors that deprive ER of its ligand, and selective estrogen receptor degraders (SERDs) such as fulvestrant are ET modalities that have been used to treat HR+ breast cancers for years (4). While, ET is the mainstay of treatment for early-stage HR+ breast cancers, about a third of patients with early stage breast cancer recur and develop resistance to ET and need other options. The development and approval of cyclin dependent kinases 4 and 6 inhibitors (CDK4/6i) a few years ago led to their use in conjunction with ET as first-line standard of care therapy for metastatic HR+ breast cancer (5). Despite the development of CDK4/6i, recurrence and resistance to ET remains a problem and greater than 50% of patients with HR+ MBC develop resistance (6). Thus, understanding the mechanisms of resistance to ET is needed to identify novel targets for patients who develop resistance. Several groups have generated *in vitro* acquired tamoxifen resistant ER+ cells lines to understand mechanisms of resistance to tamoxifen and test new therapies. Resistance to tamoxifen includes both ER dependent and independent mechanisms (7). Due to the role of growth factor receptor signaling in ET resistance, the role of growth factors including IGF1R, FGFR and ErbB2 in ET resistance have been studied (8, 9).

The type I insulin-like growth factor receptor drives proliferation, metabolism and metastasis of breast cancers (10). Insulin receptor (IR) is also overexpressed in breast cancer (11). The insulin receptor exists in two isoforms, the fetal or A isoform (IR-A) and adult or B isoform (IR-B) that are generated by alternate splicing of exon 11. IR-A lacks exon 11 while IR-B includes exon 11 which results in 12 amino acids near the juxta membrane region of the extracellular domain of IR (12). There is increased expression of IR-A in cancers compared to adult insulin target tissues (13). In liver, fat, and muscle, IR-B, serves as the receptor isoform maintaining glucose homeostasis (14, 15).

Multiple studies have shown that ERα can enhance IGF1R signaling through transcriptional upregulation of IGF1R, IRS-1 (16, 17), IGF-I (18) and IGF-II (19). IGF1R has been shown to phosphorylate and activate ER on serine-167 through a S6-kinase mechanism (20). Based on this extensive cross talk between IGF1R and ER as well as preclinical testing that showed inhibition of IGF1R inhibits HR+ tumor growth, drugs specific for IGF1R were previously tested in clinical trials in HR+ patients. However, these clinical trials did not show benefit and the addition of IGF1R antibody to ET in advanced HR+ MBC patients who had been treated with multiple endocrine therapies resulted in worse PFS (21) or did not show any added benefit (22). Retrospective analyses of archived human tumors patients treated with tamoxifen who developed resistance showed that IGF1R expression was markedly decreased (23). To further understand the clinical trial results, we previously generated *in vitro* acquired tamoxifen resistant HR+ models by culturing HR+ cells in 4-hydroxy tamoxifen (4-OHT) and showed that *in vitro* acquired tamoxifen resistant cells lose IGF1R when maintained in tamoxifen (24). These *in vitro* acquired resistant cells recover IGF1R when cultured without tamoxifen. However, due to a paucity of in vivo tamoxifen resistant models, most of the preclinical testing of IGF1R antibodies was done in endocrine sensitive models (25–27). Yet all of the clinical trials with these antibodies in breast cancer were undertaken in endocrine therapy resistant HR+ patients. It turned out one reason these trials showed lack of benefit was that these patients lacked IGF1R. and patients were not enrolled based on presence of IGF1R. Further analyses of patients in a clinical trial with IGF1R antibody revealed that disruption of IGF1R in patients caused upregulation of growth hormone and increased insulin levels (28). Genomic sequencing of a HR+ lobular breast cancer patient showed the IR gene was amplified during evolution of that patient’s tumor to endocrine resistance (29). There are some previous studies of MCF tumors in mice treated with tamoxifen that reported increased phosphorylation of IGF1R (30). It is possible that in that study it was phosphorylated IR that was detected as the phospho specific antibody used does not distinguish between phosphorylated IGF1R and IR. Thus, complete inhibition of IGF signaling may require inhibition of IR too.

The overall aim of our study was to understand if mechanisms of *in vivo* resistance to tamoxifen differed from our previously reported *in vitro* resistance by investigating the role of IGF1R and insulin receptor in *in vivo* tamoxifen resistance. Since there are not many models of tamoxifen resistance developed in vivo, we generated a new *in vivo* tamoxifen resistance model. Herein, we grew tamoxifen resistant tumors in mice until they displayed resistance *in vivo*, and established an *in vivo* tamoxifen resistant cell line. We show that *in vivo* resistance to tamoxifen is associated with irreversible loss of IGF1R. Further, in vivo tamoxifen resistance was associated with increased levels of IR, in particular the A isoform of IR (IR-A) and increased sensitivity to insulin compared to the *in vitro* acquired tamoxifen resistant cells. In vivo tamoxifen resistant cells differ from in vitro tamoxifen resistant cells and are more sensitive to insulin mediated growth. Our data suggest endocrine resistant HR+ patients may require effective inhibition of IR signaling along with the SERD fulvestrant and CDK4/6i or inhibitors of pathways implicated in resistance such as newer PI3K and Akt inhibitors.

## Materials and Methods

### Reagents

All reagents and chemicals were purchased from Sigma (St. Louis, MO) and cell culture reagents were purchased from Invitrogen/Life Technologies (Rockville, MD) unless otherwise noted. Estradiol IE2) and 4-OHT for *in vitro* experiments were prepared in ethanol at 1 mM and 40 mM respectively and diluted in serum free medium to the required concentrations. IGF-I and IGF-II were purchased from GroPep Bioreagents (Thebarton, Australia). Human insulin was from Eli Lilly (Indianapolis, IN). qScript was from Quanta Biosciences and SYBR Green from Roche. (Pleasanton, CA) or Thermo Fisher (Carlsbad, CA). The monoclonal antibody against the β-subunit of insulin receptor (IRβ) was from Santa Cruz Biotechnology, Inc. (Santa Cruz, CA). Antibodies for the β-subunit of IGF1R (IGF1Rβ), phosphorylated IGF1R/IR, p44/p42 ERK 1/2 (total) were purchased from Cell Signaling Technology (Beverly, MA). Anti-rabbit and anti-mouse secondary antibodies conjugated to HRP were from GE Biosciences/Amersham (Piscataway, NJ). Estradiol pellets were purchased from Innovative Research of America (Sarasota, FL). Acrylamide, bis-acrylamide and prestained molecular weight markers were from Bio-Rad (Hercules, CA). Tumor Dissociation kit, human (Cat # 130-095-929) was from Miltenyi Biotec (Auburn, CA).

### Cell lines and culture

All cells were grown at 37° C in a humified atmosphere with 5% CO_2_. MCF-7L cells were provided by C. Kent Osborne (Baylor College of Medicine, Houston, TX) and were routinely maintained in improved MEM Richter’s modification medium (zinc option) supplemented with 5% fetal bovine serum (FBS) and 11.25 nM human insulin, 50 U/ml penicillin and 50 µg/ml streptomycin. MCF-7L TamR were previously described (24) and C287m1-MCF-7L TamR were established from MCF-7 TamR tumors grown in athymic mice. Both of these cell lines were maintained in phenol-red free improved MEM Richter’s modification medium (zinc option) supplemented with 5% dextran charcoal characterized serum (DCC), 11.25 nM human insulin, 50 U/ml penicillin and 50 µg/ml streptomycin, and 100 nmol/L 4-OHT.

### *In vivo* xenograft growth of MCF-7L TamR tumors

All animal care and procedures were approved by the Institutional Animal Care and Use Committee and the University of Minnesota and were in accordance with the procedures detailed in the *Guide for the Care and Use of Laboratory Animals*. Mice were purchased from Envigo (Indianapolis, IN) and housed in specific pathogen free facilities. 5 ×10^6^ MCF-7L TamR cells in 50 µL phenol-red free IMEM/mouse were implanted in the second mfp on the right of 4-5 week old female athymic mice supplemented with 0.5 mg 60-day release 17β-estradiol pellet (E2) or 2 µM E2 provided in the drinking water. Tumor growth was measured twice weekly and tumor volumes were calculated using the formula length x breadth^2^/2. Five days after implantation of cells, tamoxifen citrate in peanut oil was administered subcutaneously either at a dose of 500 µg/animal (0.05 cc of the 10 mg/mL suspension) daily Monday through Friday or 1 mg/animal (0.1 cc of the 10 mg/mL suspension) administered three times weekly (Monday, Wednesday, Friday). Since tumor growth was not discerned initially, tamoxifen was stopped until tumors started growing. When tumors were about 40 mm^3^ tamoxifen was re-initiated and E2 was withdrawn four days after that. When tumor volumes reached 1300 mm^3^, tumors were harvested and necrotic areas removed. 0.35 g of the tumor from C287m1 mouse was cut with a scalpel blade and epithelial cells were dissociated into single cell suspension with a human Tumor Dissociation kit and gentleMACS™ Dissociator following the kit’s instructions for human breast xenograft tumor. Tumor was dissociated using the gentleMACS dissociator’s human tumor-01 program three times. The single cells dissociated from the *in vivo* resistant tumor were counted to determine viability and seeded in cell culture flask. When they were established and growing with consistent viability, several vials were cryopreserved and one was sent to Johns Hopkins University’s Genetic Resources Core Facility for Short Tandem Repeat (STR) profiling and analysis for authentication using the GenePrint 10 kit (Promega) and mycoplasma testing. This in vivo acquired tamoxifen resistant cell line is called C287m1-MCF-7L TamR and was maintained in medium with 100 nM 4-OHT. Cells were used in experiments after authentication. For some experiments (fig 2 and 6) where the effect of passaging in medium with and without 4-OHT was assessed, cells were passaged with 100 nM or absence of 4-OHT for eight weeks and then used for those experiments.

### RNA isolation and quantitative reverse transcription polymerase chain reaction (qRT-PCR)

Cells were seeded at a density of 2 × 10^5^ in 6-well plates. RNA was isolated using TriPure Isolation Reagent (Roche). A total of 1 μg of RNA was reverse transcribed using the qScript cDNA synthesis reagent (Quanta Biosciences) according to the manufacturer’s protocol, and qRT-PCR was performed using the Universal SYBR Green kit (Roche) or PowerTrack SYBR Green (Thermo Fisher). Target gene expression levels were normalized to the ribosomal protein, large, P0 (RPLPO). The absolute concentration of mRNA was calculated using cycle threshold values that were derived from a standard curve and normalized to RPLPO as an internal control. For quantification of INSR and its isoforms, the 2^−ΔΔCt^ method was also used to determine relative quantification of gene expression normalized to the housekeeping gene RPLPO (31). The primers used for this study are listed in Supplemental Table S1 (Supplemental File <u>1</u>).

### Cell stimulation

Cells were plated in 60-mm dishes in regular growth medium at a density of 3 to 4 × 10^5^. Cells at 70% confluence were washed two times with phosphate-buffered saline (PBS) and serum deprived for 24 hours in serum free media (SFM) as described previously (25). For treatment, medium was replaced with SFM containing either 5 nM IGF-I, 10 nM IGF-II or various doses of insulin from 0.5 to 10 nM for 10 minutes.

### Cell lysis

Cells were washed three times with ice-cold PBS on ice and lysed with 200 μl/6 cm dish lysis buffer TNESV (50 mM Tris-Cl pH 7.4, 1% NP-40, 2 mM EDTA pH 8.0, 100 mM NaCl, 2 mM Na orthovanadate) containing 25 mM glycerophosphate, 5 mM sodium fluoride, 1 mM phenylmethylsulfonyl fluoride, 20 μg/ml leupeptin, and 20 μg/ml aprotinin, Phos-STOP, and protease inhibitor cocktail. Lysates were clarified by centrifugation at 12,000 × rpm for 25 min at 4° C. Protein concentrations were determined using the bicinchoninic acid protein assay reagent kit (Thermo Fisher).

### Immunoblotting

40 µg of cellular proteins were subjected to reducing SDS-PAGE on 8% or 10% polyacrylamide gels following the Laemmli system (32). After SDS-PAGE, proteins were transferred to nitrocellulose or PVDF membrane and immunoblotted according to manufacturer guidelines.

The antibodies used for this study are listed in Supplemental Table S2 (Supplemental File <u>1</u>). Membranes were incubated with 1:1000 dilution of phospho IGF1R /pIR antibody in 2% BSA in TBST, 1:1000 dilution IGF1Rβ, phospho-Akt (Ser473), total Akt, phospho-p44/p42 ERK 1/2 (Thr 202/Tyr 204) and total p44/p42 ERK 1/2 antibodies were used as per manufacturer’s instructions.

### Growth assay for tamoxifen resistance

Tamoxifen resistance of C287m1-MCF-7L TamR line was verified by growth measurement in the presence increasing concentrations of 4-OHT after they were cultured in the presence or without 4-OHT for eight weeks. 1 × 10^4^ cells were plated in 24-well plates. The next day, cells were treated with 0 to 1000 nM 4-OHT in 1% DCC in triplicate. 96 h later, cells were trypsinized, mixed with trypan blue and counted.

### Proliferation assays

To test if E2 or insulin stimulate growth, proliferation assays with MTT (33) were used. 1 × 10^4^ or 2 × 10^4^ cells were plated in 24-well plates in 1 mL regular growth media (n=6). 24 h later, cells were switched to serum-free medium. Cells were treated with 1 or 10 nM E2 or 0 to 5 nM insulin and growth was measured 4 or 5 days later as previously described (25).

### Statistical analysis

All experiments were performed a minimum of three times. All graphs and statistical analyses were done in GraphPad Prism v9. Statistical analyses for two conditions were performed using Student’s unpaired t-test, for three or more conditions were performed using one-way ANOVA with Tukey’s multiple comparison test or Bonferroni’s multiple comparisons test. Statistical analyses of grouped cell lines were performed using two-way ANOVA with Tukey’s multiple comparison test. Error bars represent standard error of the mean (SEM) or standard deviation of variance (SD). P valuesLJ<LJ0.05 were considered statistically significant.

## Results

### *In vivo* tamoxifen resistant tumors grow in SHrN mice in the presence of tamoxifen

MCF-7L TamR cells were implanted into the second mammary fat pad on the right side of 4-5 week old female athymic mice supplemented with 2 µM E2. Mice were treated with tamoxifen 5 days after implantation of cells. Because tumor growth was not discerned at first, tamoxifen was stopped at day 25 after implantation of cells. Tumors began growing after that and when tumors were about 40 mm^3^ (day 48 after implantation of cells), tamoxifen was re-initiated. Estradiol was withdrawn at day 52 after implantation of cells. Tamoxifen did not suppress the growth of MCF-7L TamR tumors, but instead stimulated their growth (Figure 1). This accelerated growth in the presence of the drug confirmed the *in vivo* tamoxifen resistance. Mice that were implanted with MCF-7L TamR cells but were not supplemented with E2 or not treated with tamoxifen did not grow. Tumors were then harvested once tumor volumes were about 1300 mm^3^. 0.35 mg of the tumor from mouse C287m1 (Figure 1) was cut into pieces and dissociated using the human tumor dissociation kit. Dissociated cells were plated in a 100 mm dish to establish *in vivo* tamoxifen resistant line. When tumor cells were growing and viability was consistent, multiple vials were frozen and used within 18-20 passages. This line from the *in vivo* resistant tumor is referred to as C287m1-MCF-7L TamR line. STR profiling confirmed the cells were human and matched the ATCC (using ATCC’s percent match algorithm) and DSMZ databases for MCF-7 cells with 100% match.

**Figure 1.**
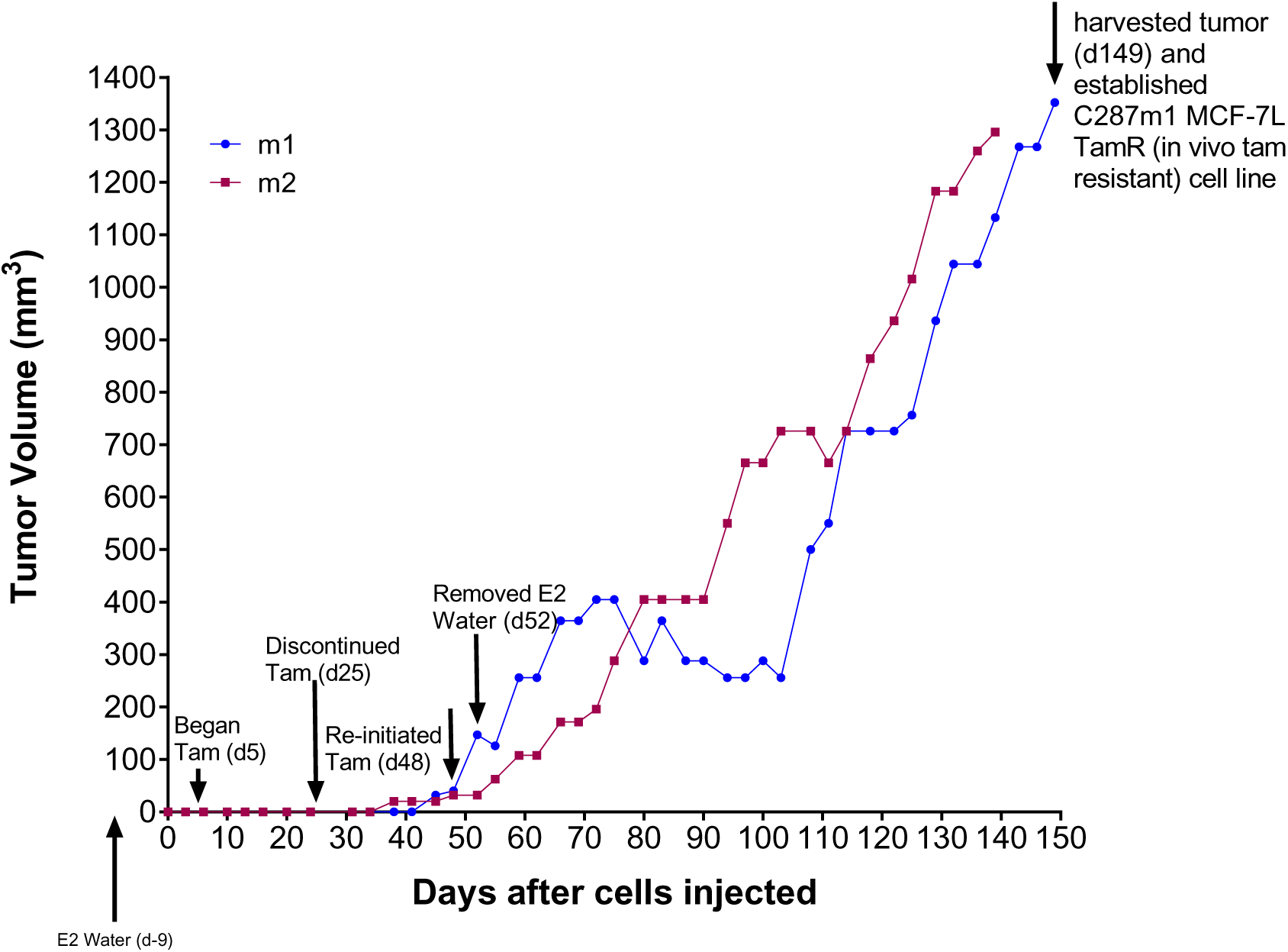
*In vivo* tamoxifen resistant tumors grow in SHrN mice in the presence of tamoxifen. MCF-7L TamR cells were implanted into the second mammary fat pad on the right of 4-5 week old female athymic mice (ShrN) supplemented with 2 µM 17β-estradiol (E2) in the drinking water. When tumors were ∼40 mm^3^, tamoxifen citrate in peanut oil was administered daily S.Q. (n = 5 mice). Tumor growth of n=2 mice is shown as tumor volumes. Tumor growth continued in the presence of tamoxifen leading to *in vivo* resistance to tamoxifen. Tumor from m1 was used to establish the *in vivo* tamoxifen resistant cells that are referred to as C287m1-MCF-7L TamR.

### *In vivo* tamoxifen resistant cells maintain resistance to tamoxifen and are growth stimulated by estrogen even when cultured in the absence of tamoxifen

To evaluate if C287m1-MCF-7L TamR cells established from the tumor retained resistance to 4-OHT *in vitro*, cells were maintained in 100 nM 4-OHT as usual or in medium without 4-OHT for 10-12 passages over 8 weeks. Cells maintained without 4-OHT were used for these experiments up to 8 additional passages so that cells cultured continuously with 4-OHT or without were used up to similar passage numbers to ensure integrity of the comparison. C287m1-MCF-7L TamR cultured with and without 4-OHT were seeded in 24-well plates in medium without any 4-OHT and treated the next day with increasing doses of 4-OHT prepared in phenol red-free medium with 1% DCC. Cell numbers were evaluated on day 4 with trypan blue. C287m1-MCF-7L TamR maintained in continued tamoxifen culture remained resistant to 1000 nM 4-0HT (Fig 2A, bars labeled +Tam) indicating resistance was maintained. Further, this resistance was maintained even when cells were cultured without 4-OHT for several passages as1000 nM 4-OHT also did not affect growth (Fig 2A, bars labeled -Tam) while we previously showed that parental MCF-7L are inhibited by that dose of 4-OHT (24). The cells maintained without 4-OHT grew slightly slower than those continuously cultured in 4-OHT. Similar to our previously described *in vitro* acquired tamoxifen resistant MC-7L TamR cells (24), E2 stimulated the growth of C287m1-MCF-7L TamR cells (Figure 2B) as measured by an MTT assay. *In vivo* tamoxifen resistant cells respond to E2 both when cultured continuously in the presence and absence of 4-OHT for eight weeks. Thus, *in vivo* TamR resistant cells remain resistant to tamoxifen and respond to E2 when cultured in the presence and absence of tamoxifen.

**Figure 2:**
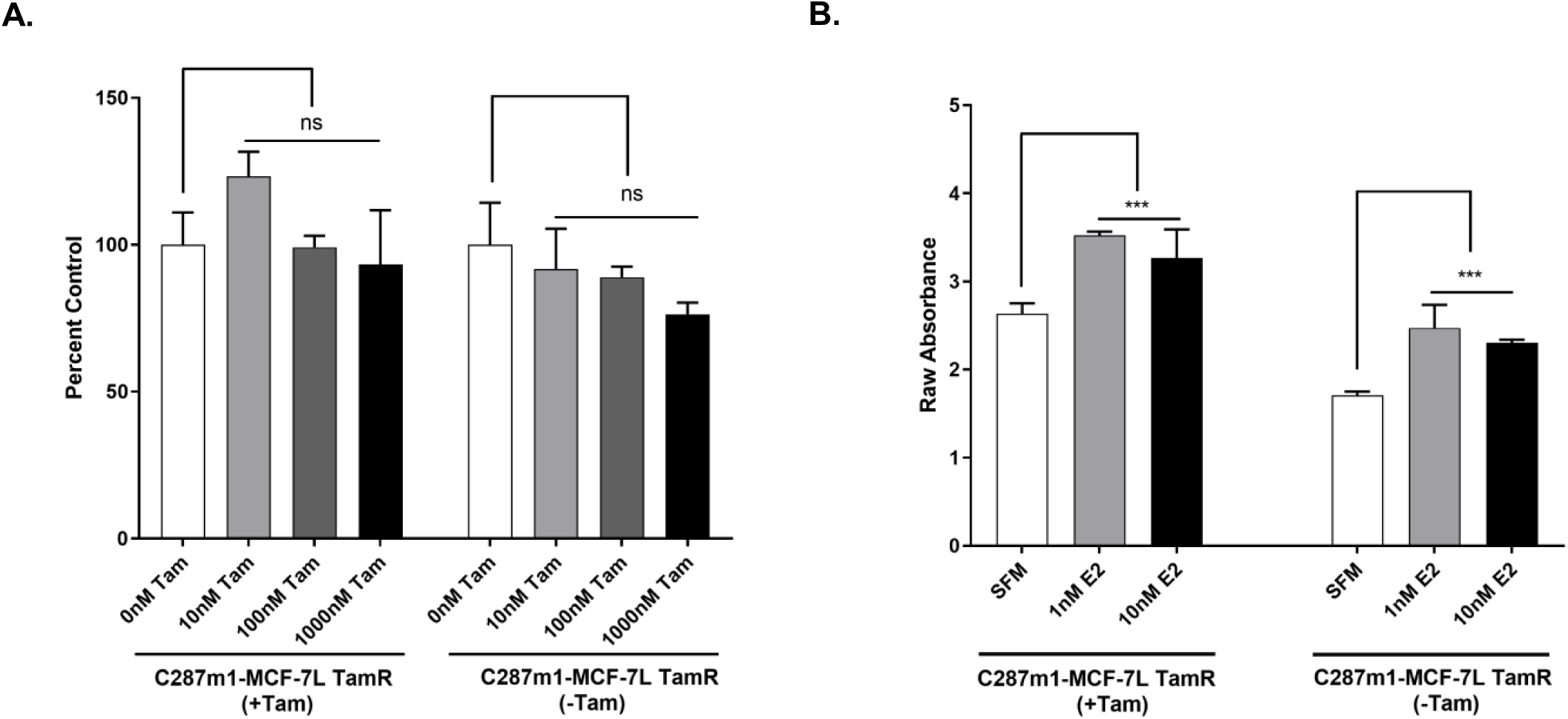
*In vivo* tamoxifen resistant cells maintain resistance to tamoxifen in vitro and are growth stimulated by estrogen even when cultured in the absence of tamoxifen. C287m1-MCF-7L TamR cells cultured continuously with 100 nM 4-OHT (graph labeled +Tam) or cultured without 4-OHT for six weeks (-Tam) were used. (**A**) Cells were plated in 24-well plates (1 × 10^4^ cells/well) in growth medium without any 4-OHT. The next day medium was replaced with phenol-red free IMEM supplemented with 1% dextran-charcoal characterized (DCC) serum and 4-OHT added at the concentrations indicated to evaluate resistance. Cell growth was measured on day 4 after staining with trypan blue and counting with a hemocytometer. Growth is shown as fold change normalized to control (0 nM 4-OHT) for each cell line. (**B**) Cells were plated in 24-well plates (1.5 × 10^4^ cells/well) in growth medium without any 4-OHT. The next day cells were starved in phenol-red free IMEM. The following day cells were treated with vehicle, 1 or 10 nM E2 in serum-free medium in triplicate and growth was measured on day 4 with MTT. Data are represented as absorbance at 570 nM. Statistical analyses were performed using two-way ANOVA with Tukey multiple comparisons test. ns = not significant, \*\*\**p* <0.001.

### *In vivo* tamoxifen resistance selectively upregulates gene expression of insulin receptor isoforms

We and others have reported that IGF1R mRNA and protein is downregulated in *in vitro* acquired tamoxifen resistance models (24, 34). Patients treated with tamoxifen also show reduced IGF1R levels (23). We next evaluated the levels of transcripts in parental tamoxifen-sensitive MCF-7L, *in vitro* acquired tamoxifen-resistant (MCF-7L TamR) and in vivo tamoxifen-resistant cells (C287m1-MCF-7L TamR). RNA was isolated from all three cell lines grown in their respective growth media. 1 µg of RNA was transcribed to cDNA and RT-qPCR with SYBR green was used to quantify transcripts. As expected, *ESR1* levels did not change across the three lines (Fig 3A left panel). *IGF1R* transcript was downregulated in both MCF-7L TamR and C287m1-MCF-7L TamR cells compared to the parental MCF-7L cells. *IRS1* levels also decreased in both the *in vitro* and *in vivo* tamoxifen resistant cells. *IRS1* encodes insulin receptor substrate 1 (IRS1) protein which is an adaptor used by phosphorylated IGF1R and IR to activate downstream signaling pathways.

**Figure 3.**
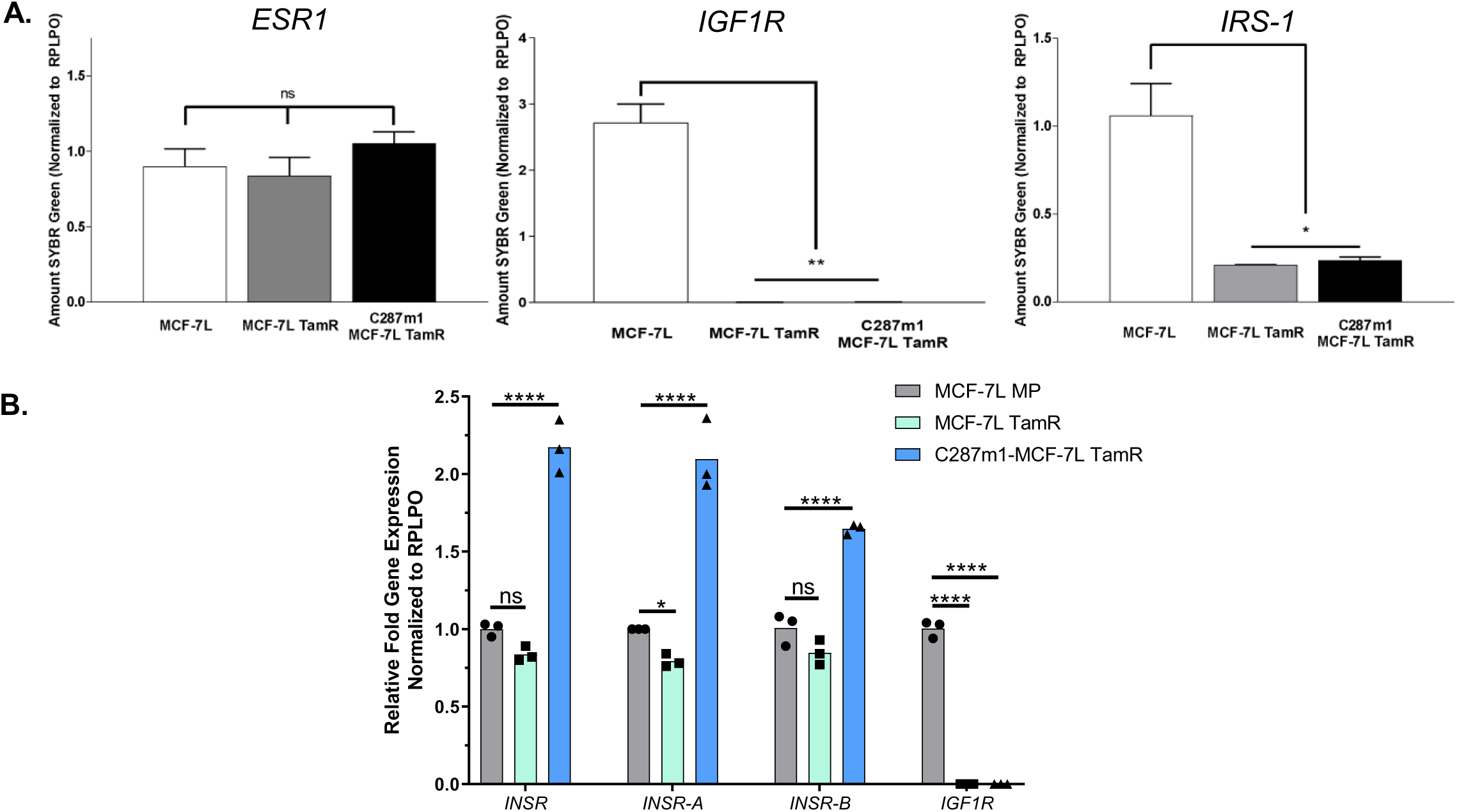
*In vivo* tamoxifen resistance selectively upregulates gene expression of insulin receptor A and B isoforms. RT-qPCR analysis was used to assess the expression of genes including: *ESR1*, *IGF1R*, *IRS-1* **(A)**, *INSR*, *INSR-A* and *INSR-B* **(B)** in parental MCF-7L (MCF-7L), *in vitro* acquired tamoxifen resistant (MCF-7L TamR) and *in vivo* tamoxifen resistant (C287m1-MCF-7L TamR) cells. Gene expression levels were normalized to levels of *RPLPO* and are shown as relative levels quantified from standard curve **(A)** and fold change over parental cell line **(B)**. Statistical analyses were performed using one-way ANOVA with Bonferroni’s multiple comparisons test **(A)** and using two-way ANOVA with Tukey multiple comparisons test **(B)**. ns = not significant, \**p* <0.05, \*\**p* <0.01, \*\*\*\**p* <0.0001.

We then checked transcript levels of total *INSR* and its two isoforms, *INSR-A* and *INSR-B. INSR-A* and *INSR-B* were detected using different forward primers designed against exon/exon junction unique to each isoform and the same reverse primer. *In vitro* acquired resistance was not associated with changes in total *INSR* or either of the isoforms (Fig 3B). These levels were slightly lower than in parental cells but the difference was not significant. Surprisingly, C287m1-MCF-7 TamR had significantly upregulated expression of total *INSR* and both *INSR-A* and *INSR-B* isoforms with *INSR-A* levels being higher. These data show that *in vivo* tamoxifen resistant cells have more total *INSR* and *INSR-A* transcripts than parental or *in vitro* resistant cells but other genes in the IGF/insulin system remained similar between the two resistant lines.

### *In vivo* tamoxifen resistance results in downregulation of IGF1R protein and increased phosphorylation of IR by insulin

We have previously showed that *in vitro* acquired resistance to tamoxifen is associated with loss of IGF1R protein and here we show IGF1R transcript was downregulated in the *in vivo* tamoxifen resistant cells (in Fig 3). Therefore, we next compared IGF1R protein levels in the three cell lines as well as signaling by the three ligands. Like the *in vitro* acquired tamoxifen resistant MCF-7L TamR, the *in vivo* tamoxifen resistant cells also showed loss of IGF1R protein compared to the robust levels of IGF1R in parental MCF-7L cells (Fig 4, upper panel) assessed by immunoblotting. Similar results were also observed by cell surface staining for IGF1R. In C287m1-MCF-7LTamR cells, IGF-I did not phosphorylate the receptor just like in MCF-7L TamR cells as they lack IGF1R. But IGF-I phosphorylated IGF1R in parental MCF-7L cells (Fig 4, lower panel) as expected and reported previously. This is in concordance with the data shown that like the MCF-7L TamR cells, C287m1-MCF-7L TamR have downregulated IGF1R compared to parental MCF-7L cells. 10 nM IGF-II phosphorylated IR in both the *in vitro* and *in vivo* TamR resistant cells but the phosphorylation was decreased compared to parental MCF-7L cells. 10 nM insulin causes enhanced phosphorylation in C287m1-MCF-7L TamR compared to the parental cells or the *in vitro* acquired tamoxifen resistant MCF-7L TamR (Fig 4, lower panel).

**Figure 4.**
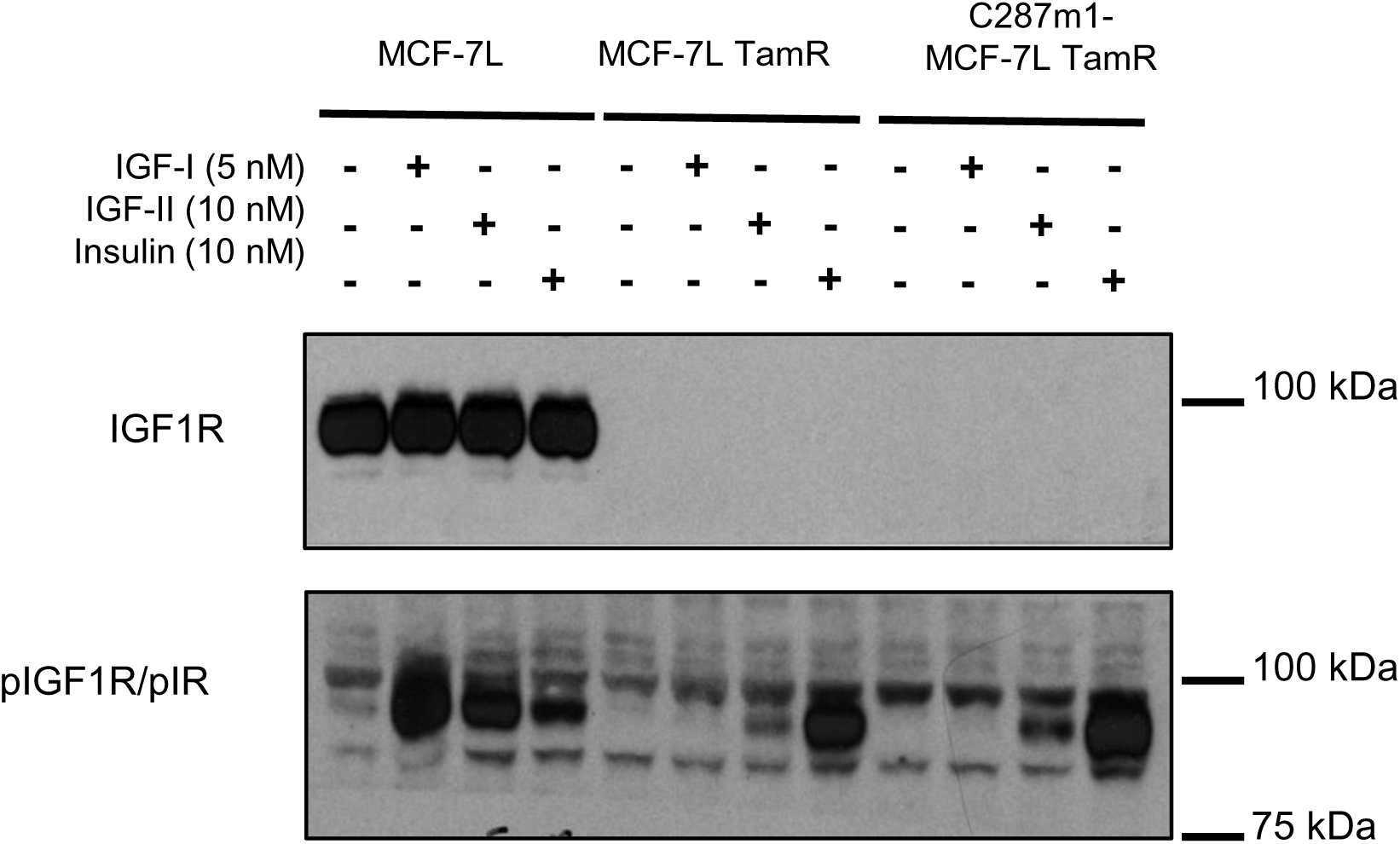
*In vivo* tamoxifen resistance results in downregulation of IGF1R protein and increased phosphorylation of IR by insulin. MCF-7L, MCF-7L TamR and C287m1-MCF-7L TamR cells were plated in 60 mm dishes. Cells were serum-starved overnight and treated for 10 minutes with 5 nM IGF-I. 10 nM IGF-II and 10 nM Ins as indicated. Cellular proteins were analyzed for IGF1R and signaling pathways by immunoblotting. The data shown are representative of three independent experiments.

### *In vivo tamoxifen* resistant cells have irreversible downregulation of IGF1R protein levels

The IGF1R protein levels recovered when the *in vitro* acquired tamoxifen resistant cells were cultured without tamoxifen and this recovery was observed in 2 to 3 weeks of culture without 4-OHT. In contrast, the loss of IGF1R protein persisted when the *in vivo* tamoxifen resistant cells were cultured without tamoxifen for 8 weeks (Fig 5). 10 minute treatment with ligands did not affect this irreversible loss of IGF1R. Thus, *in vivo* resistance is associated with irreversible loss of IGF1R protein levels. This suggests that *in vitro* acquired resistance versus *in vivo* resistance to tamoxifen results in downregulation of IGF1R through different mechanisms.

**Figure 5:**
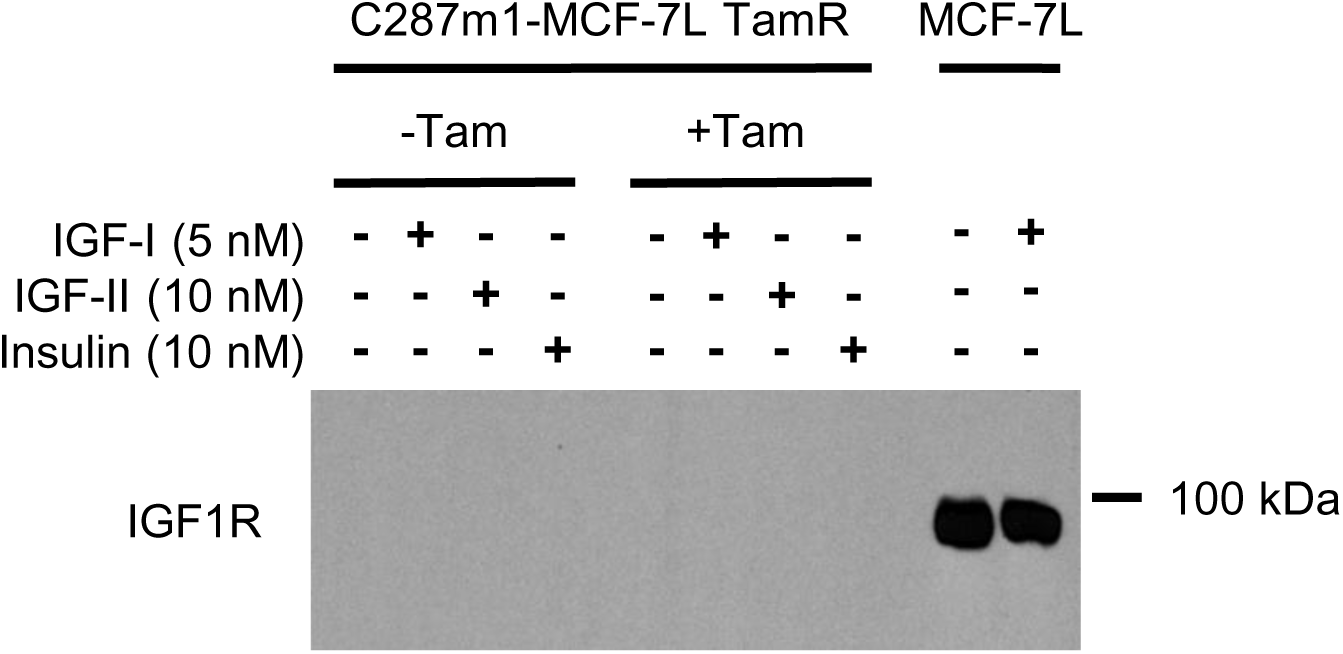
*In vivo* tamoxifen resistant cells have irreversible downregulation of IGF1R protein levels. C287m1-MCF-7L TamR cells were passaged in medium with 100 nM 4-OHT or without 4-OHT for 8 weeks. Following that time, cell lysates were collected and cellular proteins were analyzed for IGF1R by immunoblotting. IGF1R levels in control parental MCF-7L cells are shown in the last two lanes as positive controls for immunoblotting.

### *In vivo* tamoxifen resistance is associated with increased IR protein levels and enhanced sensitivity to lower doses of insulin

As *INSR* transcripts were increased and 10 nM insulin caused increased phosphorylation of IR in the *in vivo* tamoxifen resistant cells, we next examined IR protein levels in the parental MCF-7L, MCF-7L TamR and C287m1-MCF-7L TamR cells. Total IR protein levels were upregulated in C287m1 TamR compared to parental MCF-7L and MCF-7L TamR (Fig 6). Treatments with ligands for 10 minutes did not affect this upregulation. The increased IR levels in C287m1-MCF-7L TamR also caused enhanced phosphorylation of IR by lower doses of insulin than seen in the parental MCF-7L and MCF-7 TamR cells. Both 1, 2, 5 and 10 nM insulin stimulated marked phosphorylation in C287m1-MCF-7L TamR. compared to MCF-7L TamR. Only 10 nM insulin stimulated detectable phosphorylation of IR in the parental MCF-7L cells. Thus, C287m1 MCF-7L TamR have increased sensitivity to signaling by lower doses of insulin compared to parental MCF-7L and MCF-7L TamR cells with much lower concentrations of insulin (1, 2 and 5 nM) leading to enhanced phosphorylation of the receptor.

**Figure 6.**
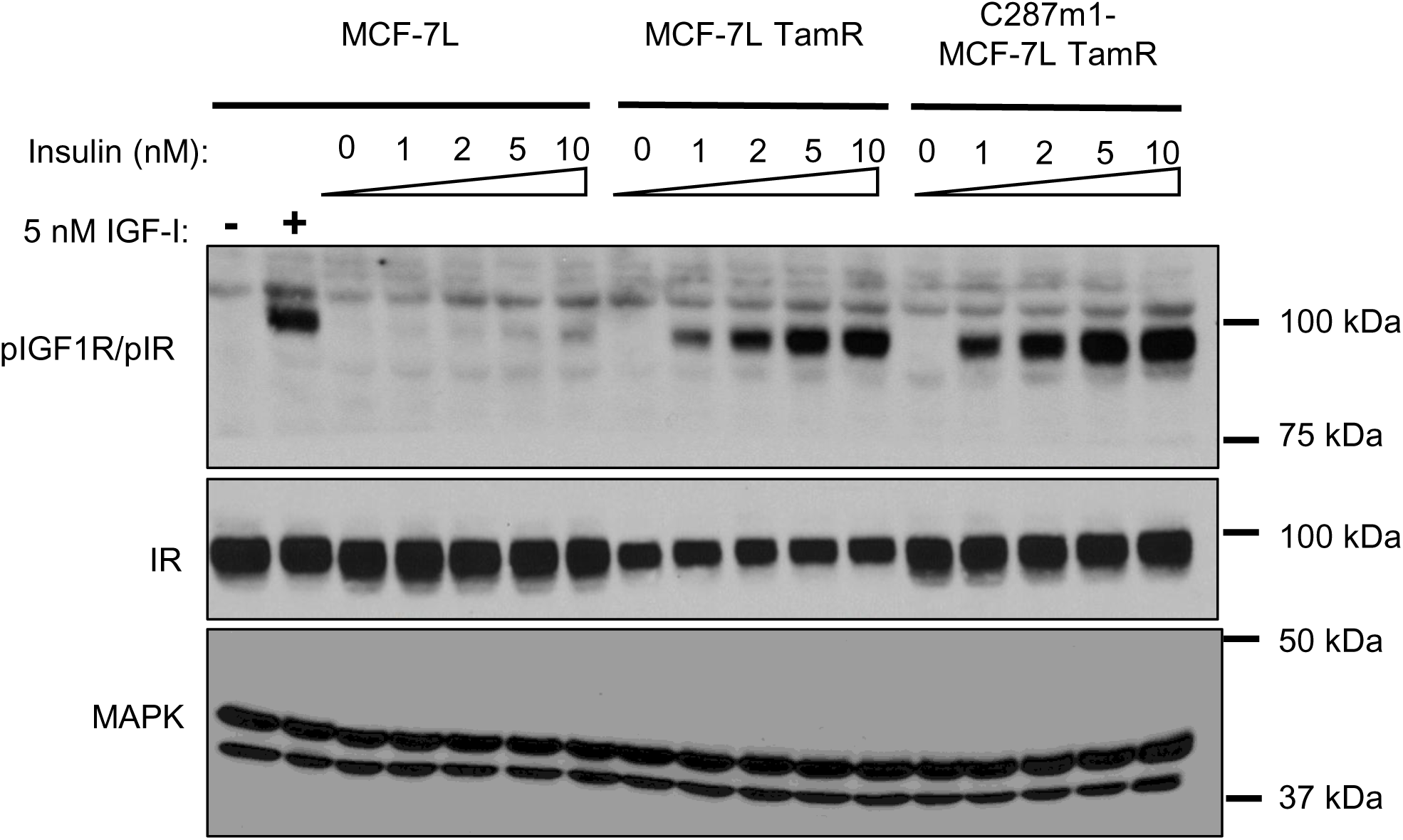
*In vivo* tamoxifen resistance is associated with increased IR protein levels and enhanced sensitivity to lower doses of insulin. MCF-7L, MCF-7L TamR and C287m1-MCF-7L TamR cells were plated in 60 mm dishes. Cells were serum-starved overnight and treated with vehicle or varying doses of insulin for 10 minutes as indicated. Cellular proteins were analyzed by immunoblotting for phosphoIR (upper panel) and total IR levels (middle panel). MAPK (lower panel) was used as loading control. The data shown are representative of three independent experiments.

### *In vivo* tamoxifen resistant cells are growth stimulated by much lower doses of insulin

Lower insulin doses also stimulated growth of C287m1 compared to parental MCF-7L and MCF-7L TamR (Fig 7). 0.1 to 2 mM insulin stimulated growth of C287m1 but not parental MCF-7L or MCF-7L TamR. In contrast, 10 nM insulin stimulated growth of all three cell lines. C287m1 MCF-7L TamR are exquisitely sensitive to growth mediated by very low doses of insulin (0,05 to 2 nM) compared to MCF-7L and MCF-7L TamR. *In vivo* tamoxifen resistant cells are growth stimulated by much lower doses of insulin compared to parental MCF-7L cells and *in vitro* acquired tamoxifen resistant cells. These data suggest that increased IR levels in the *in vivo* tamoxifen resistant cells mediates growth in response to very low insulin doses.

**Figure 7.**
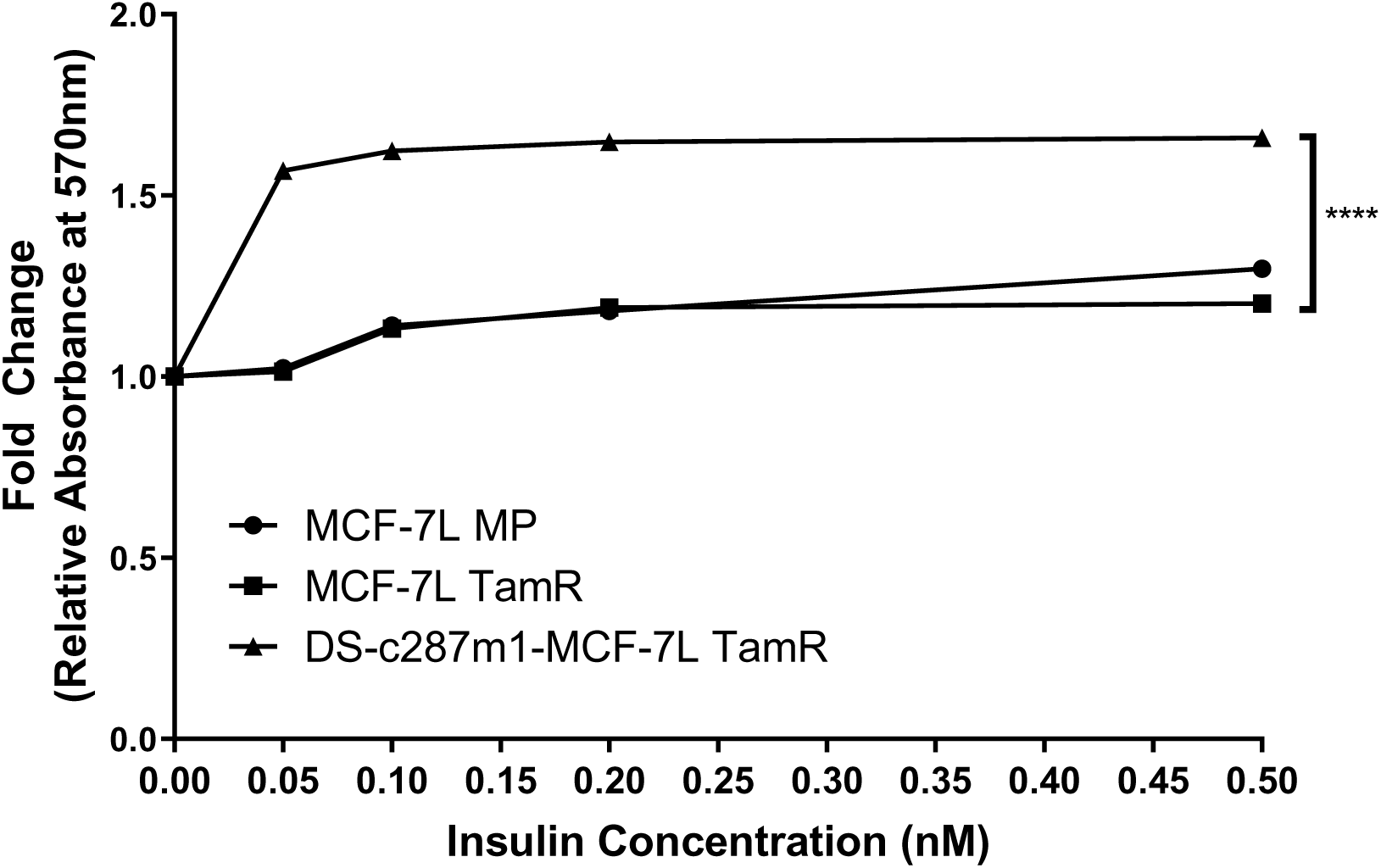
*In vivo* tamoxifen resistant cells are growth stimulated by much lower doses of insulin. MCF-7L, MCF-7L TamR and C287m1-MCF-7L TamR cells were plated in 24-well plates (1 × 10^4^ cells/well) in their growth media. Cells were serum-starved overnight and treated with varying low doses of insulin as indicated. Proliferation was measured 4 days later. Graphs represent fold change in absorbance normalized to 0 nM insulin control for each cell line. The data shown are representative of four independent experiments. Statistical analyses on absorbances were performed using two-Way ANOVA with Tukey multiple comparisons test. \*\*\*\**p* <0.0001.

## Discussion

Targeting IGF1R in clinical trials highlighted the need to use models that better reflect the clinical issue. The results of these trials showed that patients treated with IGF1R antibodies in combination with ET did worse than those treated with ET only (21, 22). All of the trials in HR+ breast cancer patients were conducted on HR+ MBC patients who had progressed on ET and had received one or multiple rounds of prior ET. Yet all of the preclinical testing was done in endocrine therapy sensitive models where combination of IGF1R antibodies and tamoxifen were more effective in inhibiting tumor growth than either alone (35, 36). The reason for this was that there are not many models of *in vivo* tamoxifen resistance. These clinical trials clearly demonstrated the need for more appropriate models to study ET resistance. There are numerous mechanisms of resistance to tamoxifen that have been investigated and reported or proposed, both ER dependent and independent (3). Our current study focused on developing an *in vivo* tamoxifen resistance model to further elucidate the role of IGF1R and IR in tamoxifen resistance due to two reasons. First, given the data from clinical trials with IGF1R antibodies showed no evidence of benefit for the combination of IGF1R antibody ganitumab with ET in resistant patients; rather the outcomes were inferior in the patients who received ganitumab with ET compared to those that received ET only (21). Second, it has been previously reported that inhibition of IR attenuated growth of tamoxifen resistant cells while the parental line that expresses both IGF1R and IR was not particularly sensitive to IR inhibition (37). Given the paucity of *in vivo* tamoxifen resistance models, here we developed a new model of *in vivo* tamoxifen resistant cells by growing *in vitro* tamoxifen resistant MCF-7L TamR tumors in mice and allowing tamoxifen resistant growth of tumors in the mice before establishing the model of *in vivo* resistant cells descried here. The *in vivo* tamoxifen resistant line maintained resistance to 4-OHT even when cultured *in vitro* without selection pressure of 4-OHT. Thus, we successfully established an *in vivo* tamoxifen resistant HR+ model.

We show that the *in vivo* tamoxifen resistant cells lose IGF1R (Fig 4). This result was expected given that we previously generated *in vitro* acquired tamoxifen resistant HR+ cell line models and reported that *in vitro* resistance resulted in reversible loss of IGF1R (24). This loss of IGF1R in *in vitro* tamoxifen resistance was later also reported by another group (34). Furthermore, following the results of the clinical trials with IGF1R antibodies combined with ER, Drury and colleagues analyzed IGF1R expression in their cohort of patients treated with tamoxifen and showed decreased IGF1R in recurrent tamoxifen resistant tumors (23). It is likely that several of the patients who were in the clinical trials with IGF1R antibodies in combination with ET had no or decreased IGF1R when they were enrolled in the trials and hence the addition of IGF1R antibody to ET in such patients showed no clinical benefit. Surprisingly, we show here that the IGF1R protein levels stay suppressed in the *in vivo* tamoxifen resistant model even when cultured in the absence of 4-OHT for eight weeks (shown in Fig 5). Thus, *in vivo* resistance results in irreversible loss of IGF1R and this loss is maintained when selective pressure of 4-OHT is removed. This differs from our *in vitro* resistance model which is associated with reversible loss of IGF1R where IGF1R protein recovers when they are cultured without the selection pressure of 4-OHT for a couple weeks. This difference in the loss of IGF1R may be because the activity of ER is differently regulated in the *in vitro* resistant versus *in vivo* resistant lines or that the mechanisms by which IGF1R is downregulated are different during *in vitro* and *in vivo* selection for tamoxifen resistance. It is also possible that the longer duration of time to reach resistance *in vivo* resulted in an irreversible loss of IGF1R as others have reported that ER levels do not recover in long term fulvestrant resistant MCF-7 while ER levels recover in short term fulvestrant resistant MCF-7 (38). While the mechanisms leading to this irreversible loss of IGF1R in the *in vivo* resistant model described here are not yet known, it provides a model that resembles what is seen in patients where IGF1R loss persists in patients who develop resistance to tamoxifen and recur.

We also show that *in vivo* resistance was associated with upregulation of *INSR*, *INSR-A* as well as *INSR-B* gene expression with *INSR-A* being more expressed. Further, we show that upregulation of *INSR*, *IR-A* and *INSR-B* transcripts was not seen in the *in vitro* tamoxifen resistant model. *INSR-A* mRNA upregulation was also seen in a clinical trial with IGF1R antibody cixutumumab with aromatase inhibitor in patients who were resistant to first-line ET. (22). Thus, the *in vivo* resistant model mimics the upregulation of *INSR-A* transcript reported in patients in this clinical trial. This trial also showed *INSR* transcript exceeded the level of *IGF1R* which is also what our study here shows in the *in vivo* tamoxifen resistant cells. Further, the *in vivo* acquired tamoxifen resistant line has significantly higher total IR protein levels detected by western blotting and flow cytometry. This increase in total IR protein levels also resulted in increased sensitivity to insulin stimulated signaling via IR as well as insulin stimulated growth. Given that the ligand IGF-II can bind to IR-A and drive tumor growth (39), this finding is significant and suggests IR-A is driving growth of the *in vivo* tamoxifen resistant cells where there is irreversible loss of IGF1R. A caveat of our study is that we were not able to measure the IR-A and IR-B protein levels due to lack of tools that specifically detect IR-A versus IR-B proteins but our study shows increased levels of total IR protein.

As there are currently only limited models to further examine the efficacy of drugs that target IR or more specifically IR-A in ET resistance, this model could potentially be an invaluable tool. This model will allow the testing of drugs currently being developed that are specific for IR-A as well as testing other drugs that target both isoforms for potential cancer therapy. Recently, several compounds have been developed such as small engineered IR-A specific binders (40) and AKS-130 (41) that can be tested using this novel *in vivo* tamoxifen resistance model where IR levels are upregulated. Patients with advanced HR+ MBC that recur on ET and CDK4/6iR in the first-line setting can be treated with SERD with and without CDK4/6i as second-line treatment. Advances in our understanding of ET resistance had led to several therapeutic options that are FDA approved or under testing in clinical trials for HR+ MBC patients who have recurred on one prior line of ET (42). Newer more recently approved SERDs such as elacestrant (43) and imlunestrant (44) which are effective in inhibiting wild type and mutant ER signaling could also be tested with IR inhibitors. As the IGF1R loss persists following tamoxifen withdrawal, the *in vivo* tamoxifen resistant cells described here will also be a valuable tool for testing the newer SERDs in combination with IR antagonists without needing tamoxifen in the combination to keep IGF1R levels low in preclinical studies. Aberrant activation of PI3K/Akt/mTORC1 including PIK3CA mutations are also associated with resistance to ET. The PI3K pathway is also activated by IGF1R and IR. Therefore, IR-A specific drugs should be tested in combination with newer drugs targeting PI3K and Akt pathways such as alpelisib (45) and capivasertib (46, 47) preclinically using the *in vivo* tamoxifen resistant model described here. Given that alpelisib and capivasertib can cause hyperglycemia, it is even more critical to test any potential future combination of these two with IR-A specific drugs in appropriate models so as not to exacerbate the hyperglycemia adverse events.

Taken together our data suggest that IR-A is a driver of tamoxifen resistance *in vivo* and the *in vivo* tamoxifen resistant cells described provide an additional model to elucidate mechanisms of resistance to tamoxifen that is relevant to what is seen in patients and will allow studies testing efficacy of drug combinations for patients with tamoxifen resistance. Certainly the advances in development and use of PDX have provided unique models of HR+ breast cancers (48, 49) but given the cost and prolonged time to grow HR+PDXs, the availability of additional readily employed model of *in vivo* tamoxifen resistance described here will also be valuable to allow for quick screening of new drugs targeting IR-A before testing them in more sophisticated PDX and PDxO models.

## Author contributions

DS conceptualized this project and planned the experiments. KJH, CSB and DS prepared the figures. KJH, PP, CSB, and VTI conducted experiments, analyzed data, and discussed results with DS. DS provided funding and supervised this study. KJH and DS wrote and revised the manuscript. All authors approved the manuscript.

## Declarations

### Ethics approval and consent to participate

All animal care and procedures were approved by the University of Minnesota IACUC and were in accordance with the procedures detailed in the Guide for the Care and Use of Laboratory Animals. No human subjects were used in this study

## Notes

Financial Support: Prospect Creek Foundation (DS), and Cancer Center Support Grant P30 CA77398

### Competing Interest Statement

The authors have declared no competing interest.

## References

1. Siegel RL, Kratzer TB, Wagle NS, Sung H, Jemal A. Cancer statistics, 2026. CA: a cancer journal for clinicians. 2026;76(1):e70043. doi: 10.3322/caac.70043. PubMed PMID: 41528114; PMCID: PMC12798275.

2. Female Breast Cancer.Subtypes: Cancer Stat Facts: Surveillance, Epidemiology, and End Results Program; National Cancer Institute. : National Institutes of Health; 2021. Available from: https://seer.cancer.gov/statfacts/html/breast-subtypes.html.

3. Hanker AB, Sudhan DR, Arteaga CL. Overcoming Endocrine Resistance in Breast Cancer. Cancer cell. 2020;37(4):496–513. doi: 10.1016/j.ccell.2020.03.009. PubMed PMID: 32289273; PMCID: PMC7169993.

4. Aggelis V, Johnston SRD. Advances in Endocrine-Based Therapies for Estrogen Receptor-Positive Metastatic Breast Cancer. Drugs. 2019;79(17):1849–66. doi: 10.1007/s40265-019-01208-8. PubMed PMID: 31630379.

5. Gradishar WJ, Moran MS, Abraham J, Abramson V, Aft R, Agnese D, Allison KH, Anderson B, Bailey J, Burstein HJ, Chen N, Chew H, Dang C, Elias AD, Giordano SH, Goetz MP, Jankowitz RC, Javid SH, Krishnamurthy J, Leitch AM, Lyons J, McCloskey S, McShane M, Mortimer J, Patel SA, Rosenberger LH, Rugo HS, Santa-Maria CA, Schneider BP, Smith ML, Soliman H, Stringer-Reasor EM, Telli ML, Wei M, Wisinski KB, Yellala A, Yeung KT, Young JS, Schonfeld R, Kumar R. NCCN Guidelines(R) Insights: Breast Cancer, Version 5.2025. J Natl Compr Canc Netw. 2025;23(11):426–36. doi: 10.6004/jnccn.2025.0053. PubMed PMID: 41213254. https://www.ncbi.nlm.nih.gov/pubmed/41213254

6. Raheem F, Karikalan SA, Batalini F, El Masry A, Mina L. Metastatic ER+ Breast Cancer: Mechanisms of Resistance and Future Therapeutic Approaches. Int J Mol Sci. 2023;24(22). Epub 20231111. doi: 10.3390/ijms242216198. PubMed PMID: 38003387; PMCID: PMC10671474.

7. Hartkopf AD, Grischke EM, Brucker SY. Endocrine-Resistant Breast Cancer: Mechanisms and Treatment. Breast Care (Basel). 2020;15(4):347–54. Epub 20200729. doi: 10.1159/000508675. PubMed PMID: 32982644; PMCID: PMC7490658.

8. Gee JM, Robertson JF, Gutteridge E, Ellis IO, Pinder SE, Rubini M, Nicholson RI. Epidermal growth factor receptor/HER2/insulin-like growth factor receptor signalling and oestrogen receptor activity in clinical breast cancer. Endocrine-related cancer. 2005;12 Suppl 1:S99–s111. doi: 10.1677/erc.1.01005. PubMed PMID: 16113104.

9. Schiff R, Massarweh S, Shou J, Osborne CK. Breast cancer endocrine resistance: how growth factor signaling and estrogen receptor coregulators modulate response. Clin Cancer Res. 2003;9(1 Pt 2):447S–54S. PubMed PMID: 12538499. http://www.ncbi.nlm.nih.gov/pubmed/12538499

10. Sachdev D, Yee D. Disrupting insulin-like growth factor signaling as a potential cancer therapy. Molecular cancer therapeutics. 2007;6(1):1–12. doi: 10.1158/1535-7163.MCT-06-0080. PubMed PMID: 17237261. http://www.ncbi.nlm.nih.gov/pubmed/17237261

11. Papa V, Pezzino V, Costantino A, Belfiore A, Giuffrida D, Frittitta L, Vannelli GB, Brand R, Goldfine ID, Vigneri R. Elevated insulin receptor content in human breast cancer. The Journal of clinical investigation. 1990;86(5):1503–10. PubMed PMID: 0002243127. http://www.ncbi.nlm.nih.gov/htbin-post/Entrez/query?db=m&form=6&dopt=r&uid=0002243127

12. Escribano O, Beneit N, Rubio-Longas C, Lopez-Pastor AR, Gomez-Hernandez A. The Role of Insulin Receptor Isoforms in Diabetes and Its Metabolic and Vascular Complications. J Diabetes Res. 2017;2017:1403206. Epub 20171019. doi: 10.1155/2017/1403206. PubMed PMID: 29201918; PMCID: PMC5671728. https://www.ncbi.nlm.nih.gov/pubmed/29201918

13. Vigneri R, Goldfine ID, Frittitta L. Insulin, insulin receptors, and cancer. J Endocrinol Invest. 2016;39(12):1365–76. Epub 20160701. doi: 10.1007/s40618-016-0508-7. PubMed PMID: 27368923. https://www.ncbi.nlm.nih.gov/pubmed/27368923

14. Flannery CA, Rowzee AM, Choe GH, Saleh FL, Radford CC, Taylor HS, Wood TL. Development of a Quantitative PCR Assay for Detection of Human Insulin-Like Growth Factor Receptor and Insulin Receptor Isoforms. Endocrinology. 2016;157(4):1702–8. Epub 20160210. doi: 10.1210/en.2015-1698. PubMed PMID: 26862994; PMCID: PMC4816738. https://www.ncbi.nlm.nih.gov/pubmed/26862994

15. Belfiore A, Frasca F, Pandini G, Sciacca L, Vigneri R. Insulin receptor isoforms and insulin receptor/insulin-like growth factor receptor hybrids in physiology and disease. Endocrine reviews. 2009;30(6):586–623. Epub 20090914. doi: 10.1210/er.2008-0047. PubMed PMID: 19752219. https://www.ncbi.nlm.nih.gov/pubmed/19752219

16. Lee AV, Jackson JG, Gooch JL, Hilsenbeck SG, Coronado-Heinsohn E, Osborne CK, Yee D. Enhancement of insulin-like growth factor signaling in human breast cancer: estrogen regulation of insulin receptor substrate-1 expression in vitro and in vivo. Molecular endocrinology. 1999;13(5):787–96

17. Molloy CA, May FE, Westley BR. Insulin receptor substrate-1 expression is regulated by estrogen in the MCF-7 human breast cancer cell line. The Journal of biological chemistry. 2000;275(17):12565–71.

18. Umayahara Y, Kawamori R, Watada H, Imano E, Iwama N, Morishima T, Yamasaki Y, Kajimoto Y, Kamada T. Estrogen regulation of the insulin-like growth factor I gene transcription involves an AP-1 enhancer. The Journal of biological chemistry. 1994;269(23):16433–42.

19. Lee AV, Darbre P, King RJB. Processing of insulin-like growth factor-II (IGF-II) by human breast cancer cells. Molecular and cellular endocrinology. 1994;99(2):211–20.

20. Becker MA, Ibrahim YH, Cui X, Lee AV, Yee D. The IGF pathway regulates ERalpha through a S6K1-dependent mechanism in breast cancer cells. Molecular endocrinology. 2011;25(3):516–28. Epub 20110203. doi: 10.1210/me.2010-0373. PubMed PMID: 21292829; PMCID: PMC3045742. https://www.ncbi.nlm.nih.gov/pubmed/21292829

21. Robertson JF, Ferrero JM, Bourgeois H, Kennecke H, de Boer RH, Jacot W, McGreivy J, Suzuki S, Zhu M, McCaffery I, Loh E, Gansert JL, Kaufman PA. Ganitumab with either exemestane or fulvestrant for postmenopausal women with advanced, hormone-receptor-positive breast cancer: a randomised, controlled, double-blind, phase 2 trial. The lancet oncology. 2013;14(3):228–35. doi: 10.1016/S1470-2045(13)70026-3. PubMed PMID: 23414585. http://www.ncbi.nlm.nih.gov/pubmed/23414585

22. Gradishar WJ, Yardley DA, Layman R, Sparano JA, Chuang E, Northfelt DW, Schwartz GN, Youssoufian H, Tang S, Novosiadly R, Forest A, Nguyen TS, Cosaert J, Grebennik D, Haluska P. Clinical and Translational Results of a Phase II, Randomized Trial of an Anti-IGF-1R (Cixutumumab) in Women with Breast Cancer That Progressed on Endocrine Therapy. Clin Cancer Res. 2016;22(2):301–9. Epub 20150831. doi: 10.1158/1078-0432.CCR-15-0588. PubMed PMID: 26324738; PMCID: PMC5548297. https://www.ncbi.nlm.nih.gov/pubmed/26324738

23. Drury SC, Detre S, Leary A, Salter J, Reis-Filho J, Barbashina V, Marchio C, Lopez-Knowles E, Ghazoui Z, Habben K, Arbogast S, Johnston S, Dowsett M. Changes in breast cancer biomarkers in the IGF1R/PI3K pathway in recurrent breast cancer after tamoxifen treatment. Endocrine-related cancer. 2011;18(5):565–77. Epub 2011/07/08. doi: 10.1530/ERC-10-0046. PubMed PMID: 21734071. https://www.ncbi.nlm.nih.gov/pubmed/21734071

24. Fagan DH, Uselman RR, Sachdev D, Yee D. Acquired resistance to tamoxifen is associated with loss of the type I insulin-like growth factor receptor: implications for breast cancer treatment. Cancer research. 2012;72(13):3372–80. doi: 10.1158/0008-5472.CAN-12-0684. PubMed PMID: 22573715. http://www.ncbi.nlm.nih.gov/pubmed/22573715

25. Sachdev D, Li SL, Hartell JS, Fujita-Yamaguchi Y, Miller JS, Yee D. A chimeric humanized single-chain antibody against the type I insulin-like growth factor (IGF) receptor renders breast cancer cells refractory to the mitogenic effects of IGF-I. Cancer research. 2003;63(3):627–35. PubMed PMID: 12566306. http://www.ncbi.nlm.nih.gov/pubmed/12566306

26. Sachdev D, Singh R, Fujita-Yamaguchi Y, Yee D. Down-regulation of insulin receptor by antibodies against the type I insulin-like growth factor receptor: implications for anti-insulin-like growth factor therapy in breast cancer. Cancer research. 2006;66(4):2391–402. doi: 10.1158/0008-5472.CAN-05-3126. PubMed PMID: 16489046. http://www.ncbi.nlm.nih.gov/pubmed/16489046

27. Burtrum D, Zhu Z, Lu D, Anderson DM, Prewett M, Pereira DS, Bassi R, Abdullah R, Hooper AT, Koo H, Jimenez X, Johnson D, Apblett R, Kussie P, Bohlen P, Witte L, Hicklin DJ, Ludwig DL. A fully human monoclonal antibody to the insulin-like growth factor I receptor blocks ligand-dependent signaling and inhibits human tumor growth in vivo. Cancer research. 2003;63(24):8912–21. PubMed PMID: 14695208. http://www.ncbi.nlm.nih.gov/entrez/query.fcgi?cmd=Retrieve&db=PubMed&dopt=Citation&list_uids=14695208

28. Haluska P, Shaw HM, Batzel GN, Yin D, Molina JR, Molife LR, Yap TA, Roberts ML, Sharma A, Gualberto A, Adjei AA, de Bono JS. Phase I dose escalation study of the anti insulin-like growth factor-I receptor monoclonal antibody CP-751,871 in patients with refractory solid tumors. Clin Cancer Res. 2007;13(19):5834–40. doi: 10.1158/1078-0432.CCR-07-1118. PubMed PMID: 17908976. http://www.ncbi.nlm.nih.gov/pubmed/17908976

29. Shah SP, Morin RD, Khattra J, Prentice L, Pugh T, Burleigh A, Delaney A, Gelmon K, Guliany R, Senz J, Steidl C, Holt RA, Jones S, Sun M, Leung G, Moore R, Severson T, Taylor GA, Teschendorff AE, Tse K, Turashvili G, Varhol R, Warren RL, Watson P, Zhao Y, Caldas C, Huntsman D, Hirst M, Marra MA, Aparicio S. Mutational evolution in a lobular breast tumour profiled at single nucleotide resolution. Nature. 2009;461(7265):809–13. doi: 10.1038/nature08489. 10.1038/nature08489

30. Massarweh S, Osborne CK, Creighton CJ, Qin L, Tsimelzon A, Huang S, Weiss H, Rimawi M, Schiff R. Tamoxifen resistance in breast tumors is driven by growth factor receptor signaling with repression of classic estrogen receptor genomic function. Cancer research. 2008;68(3):826–33. doi: 10.1158/0008-5472.CAN-07-2707. PubMed PMID: 18245484. http://www.ncbi.nlm.nih.gov/pubmed/18245484

31. Livak KJ, Schmittgen TD. Analysis of Relative Gene Expression Data Using Real-Time Quantitative PCR and the 2−ΔΔCT Method. Methods. 2001;25(4):402–8. doi: 10.1006/meth.2001.1262. https://www.sciencedirect.com/science/article/pii/S1046202301912629

32. Laemmli UK. Cleavage of structural proteins during the assembly of the head of bacteriophage T4. Nature. 1970;227:680–5.

33. Twentyman PR, Luscombe M. A study of some variables in a tetrazolium dye (MTT) based assay for cell growth and chemosensitivity. British journal of cancer. 1987;56(3):279–85. PubMed PMID: 3663476. http://www.ncbi.nlm.nih.gov/htbin-post/Entrez/query?db=m&form=6&dopt=r&uid=3663476

34. Vaziri-Gohar A, Zheng Y, Houston KD. IGF-1 Receptor Modulates FoxO1-Mediated Tamoxifen Response in Breast Cancer Cells. Molecular cancer research : MCR. 2017;15(4):489–97. Epub 20170117. doi: 10.1158/1541-7786.Mcr-16-0176. PubMed PMID: 28096479; PMCID: PMC5380564.

35. Cohen BD, Baker DA, Soderstrom C, Tkalcevic G, Rossi AM, Miller PE, Tengowski MW, Wang F, Gualberto A, Beebe JS, Moyer JD. Combination therapy enhances the inhibition of tumor growth with the fully human anti-type 1 insulin-like growth factor receptor monoclonal antibody CP-751,871. Clin Cancer Res. 2005;11(5):2063–73. doi: 10.1158/1078-0432.CCR-04-1070. PubMed PMID: 15756033. http://www.ncbi.nlm.nih.gov/pubmed/15756033

36. Ye JJ, Liang SJ, Guo N, Li SL, Wu AM, Giannini S, Sachdev D, Yee D, Brunner N, Ikle D, Fujita-Yamaguchi Y. Combined effects of tamoxifen and a chimeric humanized single chain antibody against the type I IGF receptor on breast tumor growth in vivo. Hormone and metabolic research = Hormon- und Stoffwechselforschung = Hormones et metabolisme. 2003;35(11-12):836–42. doi: 10.1055/s-2004-814145. PubMed PMID: 14710366. http://www.ncbi.nlm.nih.gov/pubmed/14710366

37. Chan JY, LaPara K, Yee D. Disruption of insulin receptor function inhibits proliferation in endocrine-resistant breast cancer cells. Oncogene. 2016;35(32):4235–43. Epub 2016/02/16. doi: 10.1038/onc.2015.488. PubMed PMID: 26876199; PMCID: PMC4982805. https://www.ncbi.nlm.nih.gov/pubmed/26876199

38. Nicholson RI, Hutcheson IR, Hiscox SE, Knowlden JM, Giles M, Barrow D, Gee JM. Growth factor signalling and resistance to selective oestrogen receptor modulators and pure anti-oestrogens: the use of anti-growth factor therapies to treat or delay endocrine resistance in breast cancer. Endocrine-related cancer. 2005;12 Suppl 1:S29–36. doi: 10.1677/erc.1.00991. PubMed PMID: 16113097. https://www.ncbi.nlm.nih.gov/pubmed/16113097

39. Sciacca L, Costantino A, Pandini G, Mineo R, Frasca F, Scalia P, Sbraccia P, Goldfine ID, Vigneri R, Belfiore A. Insulin receptor activation by IGF-II in breast cancers: evidence for a new autocrine/paracrine mechanism. Oncogene. 1999;18(15):2471–9.

40. Chan JY, Hackel BJ, Yee D. Targeting insulin receptor in breast cancer using small engineered protein scaffolds. Molecular cancer therapeutics. 2017. doi: 10.1158/1535-7163.MCT-16-0685. PubMed PMID: 28468775. https://www.ncbi.nlm.nih.gov/pubmed/28468775

41. Cao J, Yee D. Disrupting Insulin and IGF Receptor Function in Cancer. Int J Mol Sci. 2021;22(2). Epub 20210108. doi: 10.3390/ijms22020555. PubMed PMID: 33429867; PMCID: PMC7827299.

42. Tracy PD, Bopp E, Milner E, Garrido-Castro AC, Giordano A, Mayer EL, Tolaney SM, Tarantino P, Schlam I. Management of Metastatic Hormone Receptor-Positive Breast Cancer Beyond CDK4/6 Inhibitors. Current oncology reports. 2025;27(7):915–33. Epub 20250528. doi: 10.1007/s11912-025-01689-9. PubMed PMID: 40434676.

43. Bidard FC, Kaklamani VG, Neven P, Streich G, Montero AJ, Forget F, Mouret-Reynier MA, Sohn JH, Taylor D, Harnden KK, Khong H, Kocsis J, Dalenc F, Dillon PM, Babu S, Waters S, Deleu I, García Sáenz JA, Bria E, Cazzaniga M, Lu J, Aftimos P, Cortés J, Liu S, Tonini G, Laurent D, Habboubi N, Conlan MG, Bardia A. Elacestrant (oral selective estrogen receptor degrader) Versus Standard Endocrine Therapy for Estrogen Receptor-Positive, Human Epidermal Growth Factor Receptor 2-Negative Advanced Breast Cancer: Results From the Randomized Phase III EMERALD Trial. Journal of clinical oncology : official journal of the American Society of Clinical Oncology. 2022;40(28):3246–56. Epub 20220518. doi: 10.1200/jco.22.00338. PubMed PMID: 35584336; PMCID: PMC9553388.

44. Jhaveri KL, Neven P, Casalnuovo ML, Kim SB, Tokunaga E, Aftimos P, Saura C, O’Shaughnessy J, Harbeck N, Carey LA, Curigliano G, Llombart-Cussac A, Lim E, García Tinoco ML, Sohn J, Mattar A, Zhang Q, Huang CS, Hung CC, Martinez Rodriguez JL, Ruíz Borrego M, Nakamura R, Pradhan KR, Cramer von Laue C, Barrett E, Cao S, Wang XA, Smyth LM, Bidard FC. Imlunestrant with or without Abemaciclib in Advanced Breast Cancer. The New England journal of medicine. 2025;392(12):1189–202. Epub 20241211. doi: 10.1056/NEJMoa2410858. PubMed PMID: 39660834.

45. André F, Ciruelos E, Rubovszky G, Campone M, Loibl S, Rugo HS, Iwata H, Conte P, Mayer IA, Kaufman B, Yamashita T, Lu YS, Inoue K, Takahashi M, Pápai Z, Longin AS, Mills D, Wilke C, Hirawat S, Juric D. Alpelisib for PIK3CA-Mutated, Hormone Receptor-Positive Advanced Breast Cancer. The New England journal of medicine. 2019;380(20):1929–40. doi: 10.1056/NEJMoa1813904. PubMed PMID: 31091374.

46. Turner NC, Oliveira M, Howell SJ, Dalenc F, Cortes J, Gomez Moreno HL, Hu X, Jhaveri K, Krivorotko P, Loibl S, Morales Murillo S, Okera M, Park YH, Sohn J, Toi M, Tokunaga E, Yousef S, Zhukova L, de Bruin EC, Grinsted L, Schiavon G, Foxley A, Rugo HS. Capivasertib in Hormone Receptor-Positive Advanced Breast Cancer. The New England journal of medicine. 2023;388(22):2058–70. doi: 10.1056/NEJMoa2214131. PubMed PMID: 37256976; PMCID: PMC11335038.

47. Oliveira M, Rugo HS, Howell SJ, Dalenc F, Cortes J, Gomez HL, Hu X, Toi M, Jhaveri K, Krivorotko P, Loibl S, Morales Murillo S, Okera M, Nowecki Z, Park YH, Sohn JH, Tokunaga E, Yousef S, Zhukova L, Fulford M, Andrews H, Wadsworth I, D’Cruz C, Turner NC. Capivasertib and fulvestrant for patients with hormone receptor-positive, HER2-negative advanced breast cancer (CAPItello-291): patient-reported outcomes from a phase 3, randomised, double-blind, placebo-controlled trial. The lancet oncology. 2024;25(9):1231–44. doi: 10.1016/s1470-2045(24)00373-5. PubMed PMID: 39214106.

48. Dubash TD, Bardia A, Chirn B, Reeves BA, LiCausi JA, Burr R, Wittner BS, Rai S, Patel H, Bihani T, Arlt H, Bidard FC, Kaklamani VG, Aftimos P, Cortés J, Scartoni S, Fiascarelli A, Binaschi M, Habboubi N, Iafrate AJ, Toner M, Haber DA, Maheswaran S. Modeling the novel SERD elacestrant in cultured fulvestrant-refractory HR-positive breast circulating tumor cells. Breast cancer research and treatment. 2023;201(1):43–56. Epub 20230615. doi: 10.1007/s10549-023-06998-w. PubMed PMID: 37318638; PMCID: PMC10300156.

49. Guillen KP, Fujita M, Butterfield AJ, Scherer SD, Bailey MH, Chu Z, DeRose YS, Zhao L, Cortes-Sanchez E, Yang CH, Toner J, Wang G, Qiao Y, Huang X, Greenland JA, Vahrenkamp JM, Lum DH, Factor RE, Nelson EW, Matsen CB, Poretta JM, Rosenthal R, Beck AC, Buys SS, Vaklavas C, Ward JH, Jensen RL, Jones KB, Li Z, Oesterreich S, Dobrolecki LE, Pathi SS, Woo XY, Berrett KC, Wadsworth ME, Chuang JH, Lewis MT, Marth GT, Gertz J, Varley KE, Welm BE, Welm AL. A human breast cancer-derived xenograft and organoid platform for drug discovery and precision oncology. Nat Cancer. 2022;3(2):232–50. Epub 20220224. doi: 10.1038/s43018-022-00337-6. PubMed PMID: 35221336; PMCID: PMC8882468.

