## Supplemental Table for "*In vivo* acquired resistance to tamoxifen is associated with irreversible loss of IGF1R, upregulation of insulin receptor and enhanced sensitivity to insulin"

**Supplemental Table 1: qRT-PCR Primers**

| **Human Gene** | **Forward Primer Sequence (5’-3’)** | **Reverse Primer Sequence (5’-3’)** |
| --- | --- | --- |
| *ESR1* | CCC AGG GAA GCT ACT GTT TG | CTC CAC CAT GCC CTC TAC AC |
| *IGF1R* | GCA AAG GGG ACA TAA ACA CC | CAA GGC CCT TTC TCC CCA C |
| *IRS1* | TCA CAG CAG AAT GAA GAC C | CTA CTG ATG AGG AAG ATA TGA GG |
| *INSR* | CAA CGT GGT TTT CGT CCC C | AGA TGA CCA GCG ACT CCT TG |
| *INSRA* | TTT TCG TCC CCA GGC CAT C | GTC ACA TTC CCA ACA TCG CC |
| *INSRB* | CCC CAG AAA AAC CTC TTC AGG | GTC ACA TTC CCA ACA TCG CC |
| *RPLPO* | TGC TGA TGG GCA AGA ACA C | GAA CAC AAA GCC CAC ATT CC |

**Supplemental Table 2: Antibodies**

| **Antibody** | **Source** | **Cat. #** | **Dilution** |
| --- | --- | --- | --- |
| ERK 1/2 (p44/42 MAPK) | Cell Signaling | 9102 | 1:5000 |
| IGF1R β | Cell Signaling | 3027 | 1:2000 |
| IR β | Santa Cruz | sc57342 | 1:1000 |
| Phospho-IGF1Rβ (Tyr1135) and Phospho-IRβ Tyr1150 | Cell Signaling | 3918 | 1:1000 |
| Anti-rabbit IgG, HRP-linked | GE/Amersham | NA934 | 1:5000 |
| Anti-mouse IgG, HRP-linked | GE/Amersham | NA931 | 1:5000 |
